# Extracting and Interpreting the Effects of Higher Order Sequence Features on Peptide MHC Binding

**DOI:** 10.1101/2020.11.20.392233

**Authors:** Zheng Dai, Brooke D Huisman, Michael E Birnbaum, David K Gifford

## Abstract

Understanding the factors contributing to peptide MHC (pMHC) affinity is critical for the study of immune responses and the development of novel therapeutics. Developments in yeast display platforms have enabled the collection of pMHC binding data for vast libraries of peptides. However, methods for interpreting this data are still at an early stage. In this work we propose an approach for extracting peptide sequence features that affect pMHC binding from such datasets. In the process we develop the theoretical framework for fitting and interpreting these features. We demonstrate that these features accurately capture the kinetics underlying pMHC binding, and can be used to predict pMHC binding well enough to rival the current state of the art. We then analyze the extracted factors and show that they correlate with our current structural understanding of MHC molecules. Finally, we discuss the implication these factors have on the complexity of peptide engineering.

## 1 Introduction

Major Histocompatibility Complexes (MHCs) play an important role in adaptive immune responses to pathogens and cancer. Specifically, they bind peptide epitopes for recognition by T cells to facilitate T cell-mediated killing. For a peptide to be effectively presented, the strength of the peptide-MHC (pMHC) complex must be sufficiently strong. Therefore, the ability to characterize the strength of pMHC complexes is vital to understanding the immunogenic properties of peptides, and has demonstrated utility in a variety of tasks including identification of cancer neoantigens [10, 4], engineering peptides for therapeutics [15], and the selection of epitopes for vaccine inclusion [8].

Advances in deep learning have enabled the creation of predictors that are able characterize pMHC binding with high accuracy [9, 2, 6, 1, 3, 17]. However, these technologies often must wrestle with biases present in available databases, such as redundant nested peptide sets and single amino-acid variants of well-characterized peptides [11]. To extract generalizable features from training sets, these algorithms make creative use of the highly rich and complex features, which makes their trained parameters difficult to interpret [12].

Developments in yeast display platforms have enabled the unbiased collection of pMHC binding data for ~10^6^ peptides in a single experiment, and it has been shown that such data can improve existing algorithms for characterizing pMHC binding [11]. However, approaches for interpreting the data collected from these experiments is still in an early stage. In this work, we derive a principled framework for interpreting the data collected from such platforms based on reaction kinetics, and we introduce a featurization of sequences that is both sufficiently rich to model their effects on the kinetics, and is sufficiently structured for interpretation. We develop methods to extract interpretable quantities from the data and analyze the uniqueness properties of these quantities. We show that these extracted values accurately represent the reaction kinetics, and that the extracted values can be used to develop pMHC binding predictors that can rival the state of the art in the field. We find that an interpretation of our extracted features correlates with our current structural understanding of pMHC complexes. We show that it also implies the potential intractability of characterizing the set of binders for a given MHC, which carries implications for the development of peptide engineering techniques.

## 2 Interpreting the Outcome of a Peptide Display Experiment

### 2.1 Experimental Setup of the Peptide Display Assay

Here we summarize the protocol for the experimental collection of peptide-MHC binding data, and complete details can be found in [11].

The yeast display platform makes use of thousands of single polypeptide chain constructs that each encode a peptide, an antibody epitope, a cleavage site, and a class II MHC that we wish to characterize. The peptide contains a contiguous 9 residue region (the 9-mer) with high sequence diversity flanked by a set of invariant residues. The invariant residues cause binding to occur in a consistent conformation where the 9-mer is aligned to the binding pocket. This construct is expressed in yeast cells, where the peptide and the MHC will associate. The cleavage site is then cleaved and a competitor is introduced. Over a fixed period of time the peptides that bind weakly dissociate along with the antibody epitope, and the peptides that bind strongly do not. This allows enrichment of peptides that bind strongly. The entire process is then iterated over the resulting population for various rounds.

At the beginning of the experiment and after each round of enrichment, sequencing is carried out over the population to determine the distribution of 9-mers. This results in each observed 9-mer being assigned a read vector, where contained at each index *i* is the read count of that 9-mer before *i* rounds of enrichment.

### 2.2 Converting Read Vectors to Rates of Dissociation

We make the assumption that enough competitor is introduced that the amount of 9-mers that rebind to an MHC after dissociating is negligible. We model pMHC dissociation for a given peptide *s* as a first order reaction.

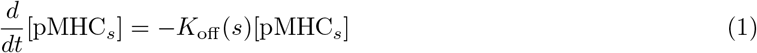

Where *K*_off_ (*s*) denotes the rate of dissociation for the 9-mer *s*, and [pMHC_*s*_] denotes the concentration of pMHC complexes between *s* and the MHC. Solving this gives the proportion of 9-mers that remain bound over the course of a round of enrichment.

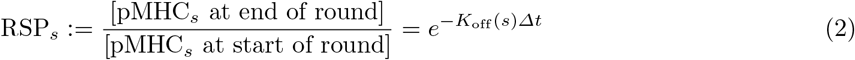

Where RSP_*s*_ is the round survival proportion or round survival probability of 9-mer *s*, and *Δt* is a fixed constant denoting the amount of time spent in one round. Since *K*_off_ (*s*) is intrinsic to *s*, so is RSP_*s*_. Let [pMHC_*s*_]_*i*_ denote the concentration MHCs complexed with *s* after *i* rounds of enrichment. We can relate the concentration between rounds with the following equation.

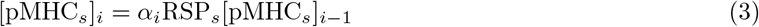

Where *α*_*i*_ is some constant that accounts for changes in the yeast cell population as a result of inter round processing. Finally, if we suppose that the read count of peptide *s* is given by a Poisson distribution with a parameter that is directly proportional to concentration, we obtain the following.

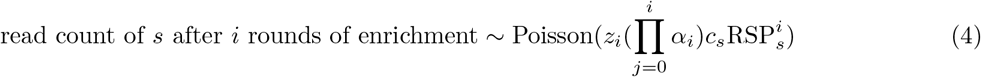

With *c_s_α*_0_ being the initial concentration of peptide *s* at the beginning of the experiment before any rounds of enrichment, and *z*_*i*_ being the constant of proportionality between concentration and sequencing reads. We assume that sequences with an all zero read vector have a known *c*_*s*_ = 0. Otherwise, we impose a prior that *ln*(*c*_*s*_) ∼ Gaussian(0, 1), which can be interpreted as modeling the initial yeast transformation and subsequent culturing as geometric Brownian motion. This prior can be seen as a regularization step.

We can then obtain an estimate of RSP_*s*_ for each peptide by solving for the maximum likelihood estimate 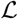 of the system.

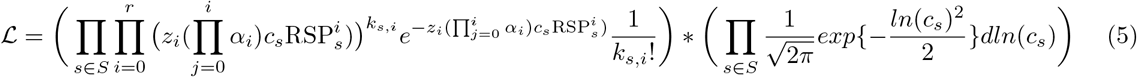

Where *S* is the set of sequences with non zero read vectors, *r* is the number of rounds, and *k*_*s,i*_ is the read count of sequence *s* after *i* rounds of enrichment. The maximum likelihood estimate is not unique, since scaling RSP_*s*_ up while scaling *α*_*i*_ or *z*_*i*_ down can give equal likelihoods. To fix this, we instead solve for *z*_*1*_*α*_0_*α*_1_RSP_*s*_ instead. We can then prove the following:

#### Theorem 1.

*Restrict* RSP_*s*_ ≥ 0 *and c*_*s*_ > 0 *for all peptides s with a nonzero read vector, and restrict α_i_* > 0 *and z_i_* > 0 *for all rounds i. Suppose that there exists at least one peptide whose read vector has no zeros at any index. Then the maximum likelihood estimate of z*_1_*α*_0_*α*_1_RSP_*s*_ for all sequences s with nonzero read vectors is unique.

This is because we can convert this into a convex optimization problem with some preprocessing. We provide the proof in appendix A. Because uniqueness conditions are easy to satisfy, going forward we will assume that they are, and we will refer to the maximum likelihood estimate of *z*_1_*α*_0_*α*_1_RSP_*s*_ as the round survival rate of *s*, denoted RSR_*s*_.

We can then rearrange equation 2 to get the following relation:

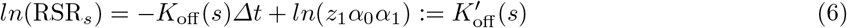

We will refer to 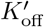 as the altered rate of dissociation for the remainder of this work.

## 3 Modelling the pMHC Dissociation Rate from Peptide Sequence Features

In this section we introduce a featurization of sequences which we will use to characterize pMHC dissociation rates. We first introduce our framework to facilitate this exposition.

### 3.1 Preliminaries

Fix *Σ* to be a finite alphabet and fix *n* to be an integer that denotes sequence length. Let [*n*] denote the set of non-negative integers that are at most *n*, and let [*n*]^+^ be [*n*] without zero. Let *Σ^n^* denote the set of all sequences of length *n*.

We define a first order sequence feature as an indicator function that operates on *Σ^n^* and outputs either a 0 or 1. A first order sequence feature is parametrized by a pair ⟨*i, c*⟩ where *i* ∈ [*n*]^+^ and *c* ∈ *Σ*, and outputs 1 on a sequence *s* if and only if *s*_*i*_ = *c*, where *s*_*i*_ is the *i*th character in *s*. We will denote such a first order sequence feature as 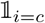, and we say 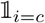 occupies position *i*. Let 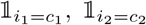 be first order sequence features. We say 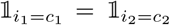 if they output the same values on all sequences. We say that 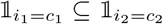 (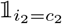 includes 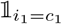) if 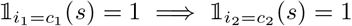 for all *s*.

We define a sequence feature generally as a product of first order sequence features that occupy distinct positions. A *k*th order sequence feature is a product of *k* first order sequence features. The first order sequence features are called the factors of the sequence feature. We extend the notions of equality and inclusion to sequence features in the same way. Let *S* be the set of positions occupied by all the first order factors of a given sequence feature. Then we say that that sequence feature occupies the positions *S*. We also introduce a unique order 0 sequence feature that outputs 1 on all sequences, denoted 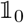, which occupies the empty set. Let *Q*_*k*_ denote a set of all sequence features of order *k*.

We define a decomposition as a mapping from sequence features to real numbers. We call the number a sequence feature maps to the coefficient of the feature, and we refer to the product of a sequence feature with its coefficient a term. We define the function of a decomposition as the sum of all its terms, and conversely we will refer to the decomposition as a decomposition of its function.

Let *D* be a decomposition. We call the restriction of *D* to only order *k* sequence features the *k*th order component of *D*, denoted *D*[*k*]. We can interpret *D*[*k*] as a decomposition itself where all sequence features that are not of order *k* are mapped to zero, so our previous definitions extend to components as well. Finally, let *k* be the largest integer such that there exists an order *k* sequence feature that *D* outputs a non-zero value on. We then call *D* an order *k* decomposition. We call a function *f* an order *k* function if there exists some *k* order decomposition whose function is *f*.

### 3.2 Rates of Dissociation are Modeled with Up to Second Order Sequence Features

For the remainder of this section with the exception of 3.4, we fix *Σ* to be the alphabet of 20 amino acids, and we fix *n* to be 9. From the previous section, we have for each 9-mer an experimentally determined 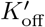 value.

Any function *f*: *Σ^n^* → ℝ can be written down as a linear combination of sequence features.

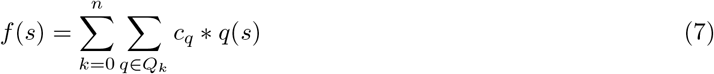

This is because each order *n* sequence feature specifies a sequence, so we can assign those coefficients the appropriate values to get a valid decomposition of *f*. For the purposes of fitting a model, this is not helpful. However, perhaps the true form of the function has a much more compact decomposition than 20^9^ coefficients. In our case, we make the following hypothesis:

#### Hypothesis

*For the purposes of modelling K*_off_, *a linear combination of sequence features up to the second order is sufficient.*

From a practical point of view, we truncate at second order because the number of third order sequence features and the number of unique 9-mers obtained from peptide display are comparable. Furthermore, not all third order sequence features are represented in the data collected from peptide display. The use of second order effects has also seen success in modelling the binding of transcription factors [13], which is an area of research that carries similarities to the one we are investigating.

Theoretically, this hypothesis can be justified from the perspective of distance based potentials, which was formulated in the context of protein structure prediction, and has seen considerable success in that field [14, 7]. In these methods, a scoring function indicating the stability of a folded protein is calculated by adding for each pair of residues a term parametrized by the identity of the pair and the distance between them. In some cases, additional terms describing the conformation are added. In the case of a pMHC complex, since 9-mers bind to class II MHCs in a conserved conformation [5], all the parameters describing conformation are fixed. The only remaining variables are then the identities of the 9-mer core, and by collecting the appropriate terms we can obtain a description in the form of a decomposition containing only second order terms.

Therefore, let 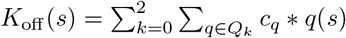. Substituting this into equation 6, we obtain:

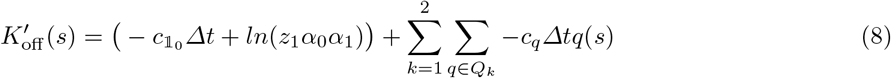

So 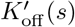 can also be described with a second order decomposition.

It is unlikely that a second order decomposition will be a perfect fit for the experimentally determined 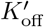 values, so we must model the error. We assume the error is primarily driven by fluctuations that randomly impact the rate of dissociation. We model this noise with a Brownian motion term *B*, and add it to equation 1 to obtain the following:

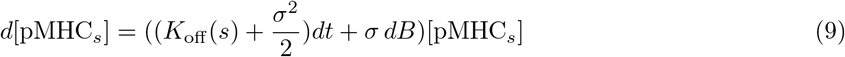

We add the 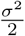 term to retain the definition of 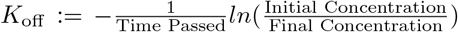 in expectation.

This is geometric Brownian motion, so integrating the above equation and substituting it into equation 6 gives the following:

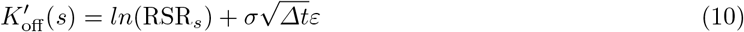

Where *ϵ* is a normally distributed random variable with zero mean and unit variance.

### 3.3 Fitting a Second Order Model to Rates of Dissociation

Since the error term in equation 10 is Gaussian, fitting a maximum likelihood model to the data is equivalent to minimizing the squared error. However, a complication arises if we assign 0 to RSR_*s*_ for any *s*, since *ln*(0) = −∞. This occurs for a substantial portion of the dataset, so instead of throwing these values away we will instead treat these as a limitation of measurement. Specifically, we assume that the smallest finite *ln*(RSR) value represents the smallest measureable value, which we will refer to as *ln*(*ϵ*). We then suppose that RSR values that equal zero actually have a finite *ln*(RSR) that is at most equal to *ln*(*ϵ*). We then assign to them values that maximizes the likelihood subject to that constraint.

To formalize, let *F*_2_ be the set of functions that can be described by a second order decomposition. Let *S*^+^ be the set of sequences that have positive RSR and *S*^0^ be the set of sequences that have zero RSR. We then want to minimize the following:

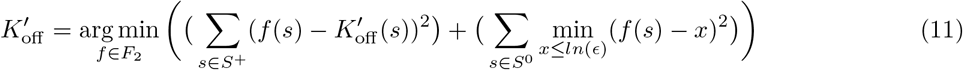

To solve this, note that functions in *F*_2_ are linear combinations of sequence features. We fix an ordered list |*L*| containing all sequence features in *Q*_0_, *Q*_1_, and *Q*_2_. We can then encode each sequence *s* as a vector *v*_*s*_ of length *L*, where we place *L*_*i*_(*s*) at index *i*, *L*_*i*_ being the *i*th element of *L*. Then any function *f* ϵ *F*_2_ can also be encoded as a vector. Let *D* be a second order decomposition of *f*. We can then encode *f* in a vector *v*_*D*_ where at index *i* we have the coefficient *D*(*L*_*i*_). We can then compute 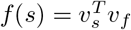.

The representation of *f* as *D* is not unique. This implies the existence of multiple solutions if we attempt to solve for a vector representation of *f*. To remove this redundancy, we fix an arbitrary symbol INV ∈ *Σ*, which we call the invariant symbol. We then remove from consideration all sequence features that contain a first order factor of the form 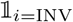.

#### Theorem 2.

*Let* 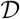 *be the set containing all decompositions D where for any sequence feature* 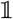 *that contains a first order factor of the form* 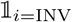, *the coefficient associated with it is* 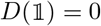. *Let f be any function. Then there exists exactly one decomposition in* 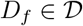 *whose function is f. Furthermore, if f is an order k function, for any sequence feature* 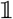 *of order greater than k*, 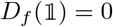.

A proof is given in appendix B. Thus, by fixing an invariant symbol and discarding all sequence features containing it, we can construct a set of vector representations of the sequences and functions as above, but in such a way that each vector uniquely defines a function. Using this representation, we can reformulate our optimization:

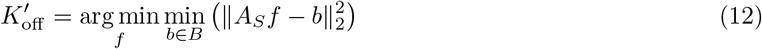

Let *S*^ord^ be an ordered list of all sequences under consideration. Then *A*_*s*_ is the matrix where the *i*th row is equal to the row vector representation of the *i*th sequence 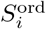. Let *B* be the set of vectors where the value at index *i* is *ln*(RSR ord) if it is finite, or some number no larger than *ln*(*ϵ*) otherwise.

Note that *B* is an intersection of half spaces, and *A_S_ f* resides within a hyperplane. The objective then boils down to minimizing the distance between a pair of convex sets. We can converge on a solution by iteratively projecting between the sets. Projecting onto the column space of *A*_*S*_ can be done by solving 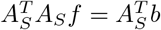, and each iteration can be performed efficiently if the LU-factorization of 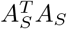 is precomputed. The most expensive step per iteration then becomes either computing 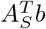 and *A_S_ f* in time 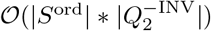, or back substituting in time 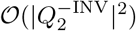, where 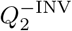 INV denotes the set of sequence features up to order 2 that don’t contain INV. Projecting onto *B* can be done componentwise since the surfaces of *B* align with the coordinate axes, so can be done in time 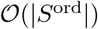 per iteration.

### 3.4 Interpreting Decompositions

Since we could have fixed any symbol as our invariant, the fact that the coefficients associated with that symbol are zero has no intrinsic meaning. In general, the coefficients of an arbitrary decomposition of a function tell us little about the structure of the function. However, there exists a canonical decomposition whose coefficients are fixed and carry meaning.

Let *S* denote a random sequence that is distributed uniformly over *Σ^n^*. A desireable property for a decomposition to have would be to maximize the “impact” of the lower order sequence features. For example, order *n* sequence features are simply corrections applied to each sequence, and says little about the structure of the function, so those should be minimized. This motivates the definition of parsimony, which formalizes this concept.

#### Definition 1.

*Let D and D’ be decompositions. Denote the function of the kth order components of D and D’ as* 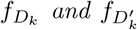 *respectively. Then D at least as parsimonious as D’ if for any k*, 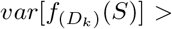 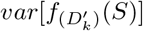 *only if there exists some k’* > *k such that* 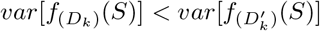.

Suppose we have a function *f*: *Σ*^*n*^ → ℝ. Then we define the canonical decomposition of *f* as the decomposition *C* with the following inductively defined coefficients:

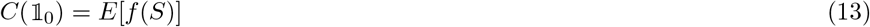

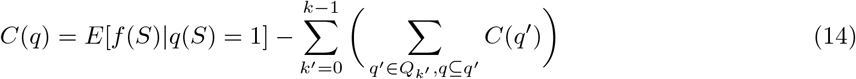

Where equation 14 defines *C*(*q*) for all order *k* sequence features *q* that are not 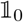 for all *k*. This decomposition is well defined since the induction can start from 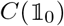, and coefficients at each subsequent order are only defined in terms of coefficients that are of lower order.

#### Theorem 3.

*Given a function f, the canonical decomposition of f is a decomposition of f and is unique. Furthermore, the canonical decomposition of f is at least as parsimonious as any other decomposition of f.*

A proof is provided in appendix C.

The canonical decomposition assigns coefficients to sequence features in a way that captures their intrinsic statistical impact on the function, and captures them at the lowest possible order. As a corollary, we note that if *f* is a *k*th order function, then because its canonical decomposition is maximally parsimonious, all sequence features whose orders are greater than *k* must be assigned zero in the canonical decomposition. Therefore, if we are already given a low order decomposition of *f*, we would know to stop at a given order. Furthermore, due to the linearity of expectation, calculating expectations is efficient relative to the size of a decomposition, so computing a canonical decomposition is efficient if we already have a low order decomposition.

## 4 Results and Discussion

### 4.1 Fitted Models Reproduce Motifs Found in Experimental Data

We apply our methodology to peptide display data collected from two class II MHCs: DR401 (HLA-DRA1*01:01, HLA-DRB1*04:01) and DR402 (HLA-DRA1*01:01, HLA-DRB1*04:02). While the two alleles are highly similar, they differ in a few key residues within their binding pockets, which shifts their binding preferences [11]. Therefore, applying our methodology to both alleles allows us to make statements about its generalizability.

Peptide display was conducted with 5 rounds of enrichment for both alleles. We fit for both alleles a second order function approximating their altered rate of dissociation 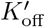, and we fix tyrosine (Y) as the invariant symbol for the fitting process. We will refer to these functions as 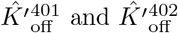 for the remainder of this manuscript. Using 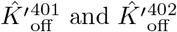, we can run an experiment analogous to the peptide display experiment *in silico*, where we start with a uniform distribution of sequences and see how the distribution shifts as time progresses. We describe this in more detail in appendix D. Since *Δt* is built into the altered dissociation constant, running for 1 unit of time is equivalent to running the experiment for 1 round, thus we can directly compare our results against the experimental results.

We show the 9-mer motifs we obtain after 5 rounds compared with the experimentally obtained motifs in figure 1. Figures showing the results after all 5 rounds are given in appendix D. Although magnitudes differ, the overall residue preferences are well represented by the simulation.

**Fig. 1.**
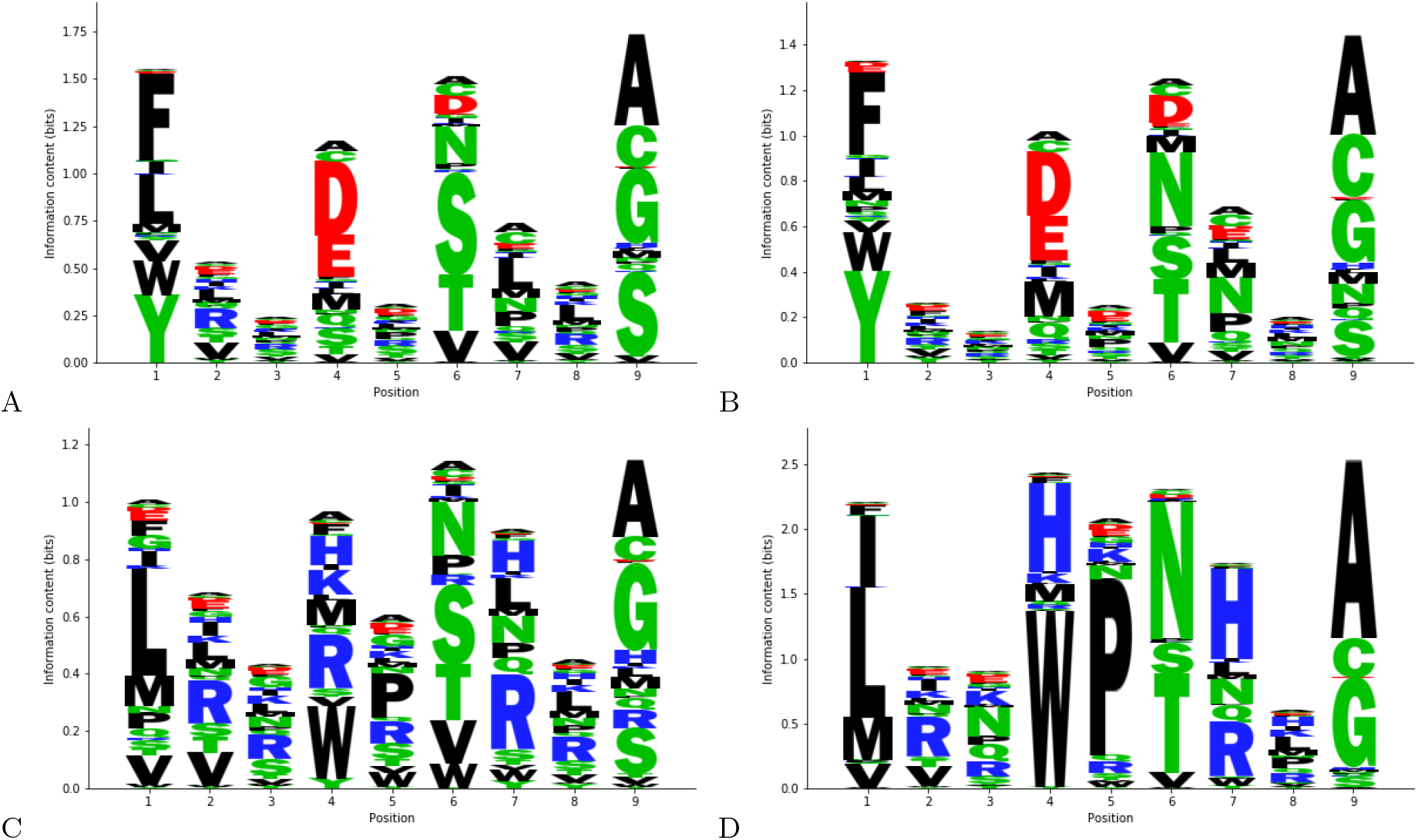
Simulated runs of peptide display reproduce binding preferences seen in experimental data. The distribution of peptide sequences obtained after running the peptide display assay *in silico* using the modelled *K*_off_ values starting with a completely uniform distribution of sequences (B,D), compared against the empirical distribution obtained from peptide display (A,C). Shown are the outcomes after round 5 depicted as sequence logos. (A,B) depict DR401, and (C,D) depict DR402.

### 4.2 Rates of Dissociation is a State-of-the-Art Predictor of Peptide Binding

Although 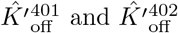 give rates of dissociation rather than the equilibrium constants more commonly used as measures of affinity, we can still characterize peptides with a high *K*_off_ as poor binders. Therefore, our model can be used as a way of predicting pMHC binding.

To test this, we benchmark 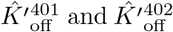 against various state-of-the-art algorithms using two datasets derived from eluted ligand monoallelic mass spectrometry (MS), one dataset derived from yeast display (YD), and one dataset generated from fluorescence polarization competition assays (FP) [11]. Since these datasets are not comprised of 9-mers, we must first identify the correct binding site. We do this by considering each contiguous 9 residue window, and taking the window with the highest altered rate of dissociation as the binding site, the rationale being that most peptides would be bound in the most preferable conformation.

The results are presented in figure 2, benchmarked against various state-of-the-art algorithms [6, 1, 3], and two deep learning architectures trained with the same yeast display data we used to extract our altered rates of dissociation [9, 17]. We see that both 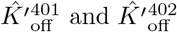 achieve highly competitive results. The yeast display models tend to perform better with YD data, while still achieving good performance in the MS datasets. The converse is not true, where the non YD trained models are significantly outperformed on the YD test set, while they tend to perform slightly better on MS datasets. We remark that both deep learning architectures are unable to significantly outperform the predictions made by altered dissociation rates, this being especially the case for DR401.

**Fig. 2.**
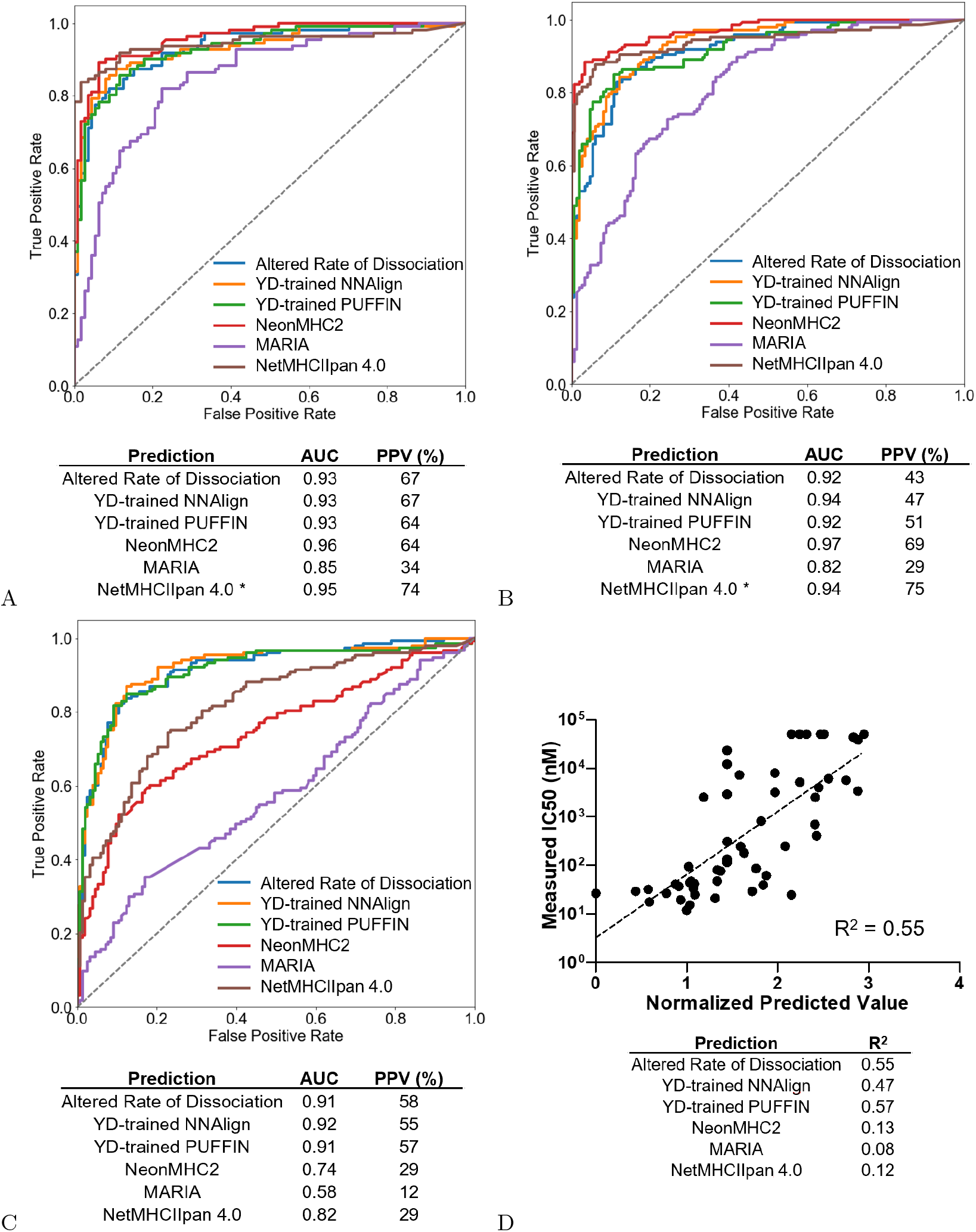
Altered rates of dissociation is a good predictor of pMHC binding. We benchmark our model (altered rate of dissociation) against various state-of-the-art prediction algorithms on four benchmark datasets used in [11]. We also benchmark against versions of NNAlign and PUFFIN that were trained on the same yeast display data used to extract the altered rates. Here we plot the auROC curves and compute the positive predictive value for each model for three of the datasets (A,B,C). An asterisk beside a model indicates that the benchmarking set intersects with the dataset used to train the model. A: contains a benchmark against DR401 data from eluted MS ligands. B: contains a benchmark against DR402 data from eluted MS ligands. C: contains a benchmark against a DR401 dataset from a different yeast display assay on randomized 13-mers that don’t have a fixed 9-mer for binding. D: compares performance by looking at how well outputs correlate with the logarithm of measured IC50 values for DR401. We report the squared correlation coefficients for each model. The normalized predicted value plotted is the altered rate of dissociation shifted so the smallest value is 0.

It appears to be the case that deep learning architectures are still able to extract features that slightly improve predictions for DR402 binding. DR402 has a truncated binding pocket in its binding grove, which may translate into smaller first order effects overall. Therefore, we hypothesize there are indeed higher order sequence features that play a role in pMHC binding. They are negligible in DR401 since they are overshadowed by the overall stronger lower order effects, but become noticeable when predicting DR402 binding.

We remark that because 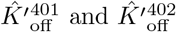 achieve high accuracy, that suggests that our methodology for selecting the binding site is reasonably accurate, and so 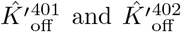 can also be used to predict the binding sites.

### 4.3 Visualizing the Factors Impacting the Rate of Dissociation

We compute the canonical decompositions of 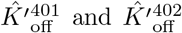, denoted *D*_401_ and *D*_402_ respectively. The structure of second order decompositions are highly complex, so to facilitate their interpretability we sum the terms related to sequence features that occupy the same position. We refer to this sum as a positional function, and each positional function is indexed by a unique set of positions. Ignoring the zeroth order positional function (which has little meaning when considering altered rates of dissociation), we can lay out the positional function in a matrix, where at each pair of distinct indices we place the positional variable indexed by those indices, and along the main diagonal we place the first order positional functions, such that the variable that occupies *i* is placed at *i, i*.

If we let *S* be a sequence drawn uniformly at random, then a positional function on that sequence is a random variable. We can then characterize the “impact” of a positional function by its variance. This is equivalent to squaring the coefficients for each term in a positional function and taking their average. We can also characterize how similar the impacts of each position are by looking at the correlation coefficient of a pair of positional functions using two independently and uniformly drawn random sequences.

We plot the variances and covariances of the positional functions in figure 3. We can immediately see that along the main diagnoal positions 1, 4, 6, and 9 have strong first order variances, which is consistent with our structural understanding of the molecules [5]. We also see that the relative first order effect of position 1 is significantly reduced in DR402 compared to DR401, which may be due to the truncated binding pocket in DR402 compared to DR402. Interestingly, it appears that this is reveals a more impactful second order interaction between position 1 and position 6, which is consistent with our earlier hypothesis that higher order effects play a less negligible role in DR402 binding preferences. We can also see that the most correlated positional function between the two alleles is at p9, which is the conserved binding pocket.

**Fig. 3.**
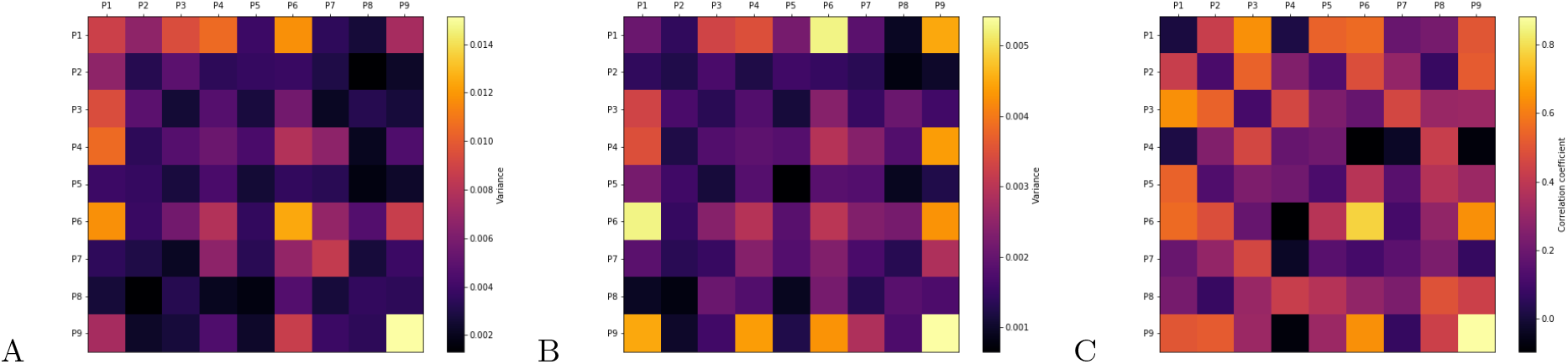
Visualizing the canonical decompositions of 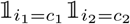. For each pair of positions we plot quantities relating to the positional function associated with that pair. Along the diagonal are the quantities relating to the positional functions associated with first order features. Note that the matrices are symmetric, since positional functions are indexed by unordered sets. A: We plot the variances of the positional functions for DR401. B: We plot the variances of the positional functions for DR402. C: We plot the correlation coefficients of the positional functions that occupy the same positions between DR401 and DR402.

### 4.4 Optimization Becomes Difficult in the Presence of Higher Order Effects

Once a function is understood, it is usually desireable to use it as an objective for optimization. For example, it has been shown that altering peptide residues to optimize MHC binding can improve the immunogenetic properties of peptides [15]. Supposing that the impact of second order terms are negligible, then certain optimization tasks become trivial. For example, we can efficiently find the best binder for a specified MHC given a first order decomposition characterizing its binding by greedily selecting the residue that has the most beneficial effect at each position. Unfortunately, the canonical decomposition pictured in figure 3 suggests that the impact of second order sequence features is comparable to the impact of first order features, and so they are vital for accurately characterizing binding. Consequentially, the task of finding the best binder for a specified MHC given a second order decomposition characterizing its binding becomes NP-hard.

#### Theorem 4.

*Given a second order decomposition of a function f: Σ^n^* → ℝ *and some k, the problem of determining whether there exists a sequence s* ∈ *Σ^n^ such that f* (*s*) > *k is NP-complete if* |*Σ*| ≥ 3.

*Proof.* We can reduce from 3-Coloring on *n* nodes. Partition elements of *Σ* into 3 sets corresponding to each color. Assign each position a distinct node. Assign zero coefficients to all first and zeroth order sequence features, and assign zero coefficients to second order sequence features 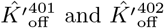 unless the nodes assigned to *i*_1_ and *i*_2_ contain an edge between them, and if *c*_1_ and *c*_2_ belong to the same color partition, in which case assign it a coefficient of −2. Assigning a residue to a position is then equivalent to assigning a color to a node, so there exists a sequence *s* such that *f* (*s*) > −1 if and only if the original graph is 3-colorable. This completes the proof.

While finding the optimal value is a specific task, the proof reveals that a second order decomposition encodes a graph structure. It is then not difficult to see how other peptide engineering tasks can be similarly reduced in this way. In our case *n* is only equal to 9. While potentially unpleasant, this is not intractable, especially with the use of heuristic speedups. However, it is known that residues flanking the 9-mer core can affect binding [16], which already suggests that future work may wish to take these effects into account. It is also likely that second order effects play important roles in more complex systems that involve longer sequences, which will necessitate the development of efficient heuristics to tackle these optimization tasks.

## 5 Conclusion

We have developed a method of extracting the reaction kinetics underlying large yeast displayed libraries. We provided a method of extracting well defined estimates of the reaction kinetics, and introduced a featurization of sequences that is able to accurately capture the factors that influence the reaction. The factors we extract are accurate to the point where we can use them to construct a pMHC binding predictor that rivals state-of-the-art deep learning architectures, many of which express a far larger parameter space.

Furthermore, we provide a framework for interpreting our featurization of sequences. We showed that this framework reveals known factors that influence pMHC binding, and we expect it can be further exploited to reveal novel insights in this domain. The framework is also highly generalizable in both the sense that it can be applied to richer sets of sequence features, and that it applies to arbitrary finite sequence valued domains, and we hope to leverage it to its full potential in the future.

## Appendix

## A Proof of Theorem 1 (Uniqueness of RSR)

We show in this section that RSR values are uniquely defined for all sequences with non zero read vectors if there exists at least one sequence whose read vector contains no zero entries. Without loss of generality, suppose there are no sequences with zero read vectors, since they play no role in the likelihood. The likelihood 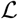 we want to optimize then takes the following form:

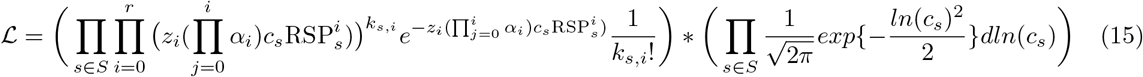

Where *S* is the set of all sequences, *r* is the number of rounds of enrichment, and *k*_*s,i*_ is the read count of sequence *s* after *i* rounds of enrichment (the *i* + 1st entry of the read vector of *s*). The first product is a product of poisson distributions, while the second project is a product of log normal distributions. To simplify the form, we introduce the following variables:

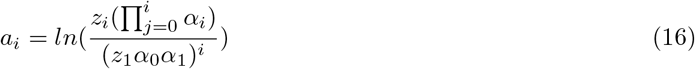

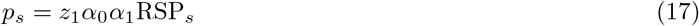

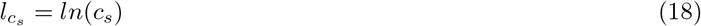

These values are always well defined the estimated parameters *α*_*i*_, *z*_*i*_, and *c*_*s*_ are restricted to be strictly positive. Note that what we are trying to show is equivalent to showing that *p*_*s*_ are uniquely defined. We can then reformulate our likelihood function:

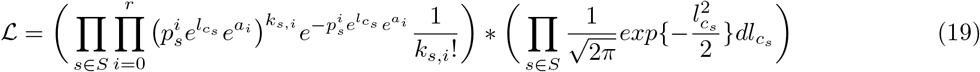

For all sequences *s* with a read vector that is all zero past the first entry, setting *p*_*s*_ = 0 clearly gives the maximum likelihood. Conversely, if there are any non zero entries past the first entry, then *p*_*s*_ cannot be zero in the maximum likelihood estimate since that gives a likelihood of zero. Let *S*^+^ denote the set of sequences whose read vector contains non zero entries past the first one. Then the maximum likelihood estimate is also the optimum of the following modified likelihood 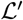:

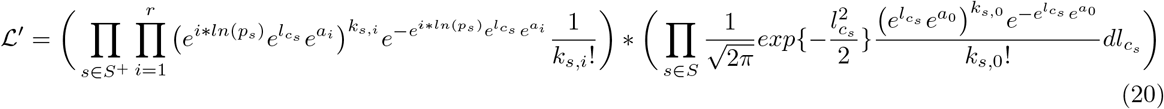

Where we moved some terms from the first product to the second, and we are allowed to write *p*_*S*_ as 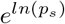 for all *s* in *S*^+^ since *p*_*s*_ is not allowed to be 0. Let 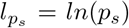. This likelihood is optimized when the log likelihood function 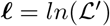 is optimized.

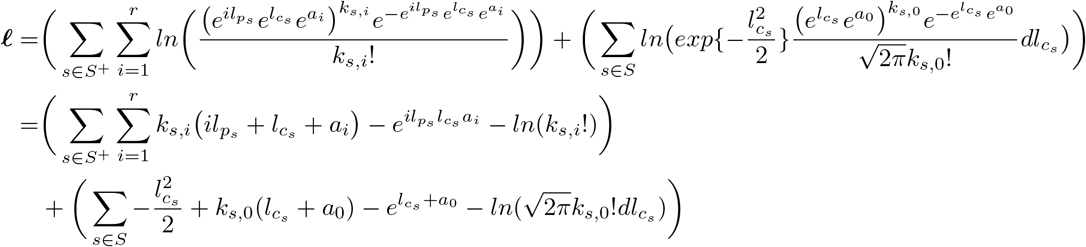

Since constant terms do not affect optimality, if we drop them then the resulting 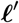 is optimized if and only if 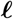 is optimized:

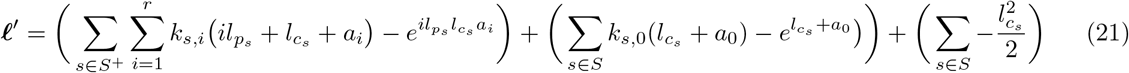

For convenience, we will define the family of functions *F*_*c*_ parametrized by a *c* ≥ 0. We will also define *G*:

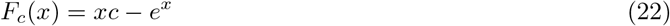

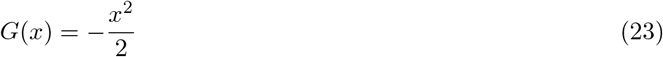

We can then rewrite equation 24 as the following:

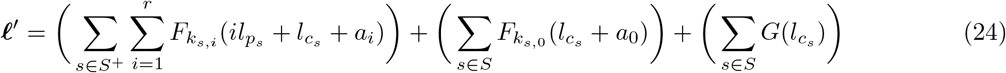

### Lemma 1.

*For all c, F_c_ and G are strictly concave and bounded from above. Furthermore, with the exception of F*_0_*, they all approach* −∞ *as their arguments approach* ∞ *or* −∞.

*Proof. G* is a parabola, so the statements follow. To see that *F*_*c*_ is strictly concave, note that the *xc* term is linear, while −*e*^*x*^ is strictly concave. Thus, their sum must be strictly concave. To see that it is bounded from above, we take the derivative 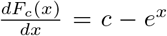, which becomes negative from sufficiently large *x*, and positive for all sufficiently small *x*. Since the derivative only tends to 0 on either extreme if *c* = 0, *F*_*c*_ drops without bound on both extremes when *c* ≠ 0.

### Lemma 2.

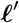 *is strictly concave*.

*Proof.* Suppose not. Then there exists some set of parameters 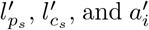, some different set of parameters 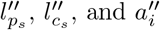, and some *x* > 0 and *y* > 0 where *x* + *y* = 1 such that 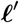 evaluates to *L*’ on the first set and *L*’ on the second set, and where given parameters 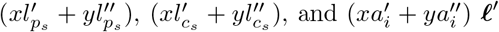 evaluates to *L*_*xy*_ ≤ *xL*’ + *yL*”.

Define 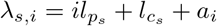 as a shorthand. Then we have:

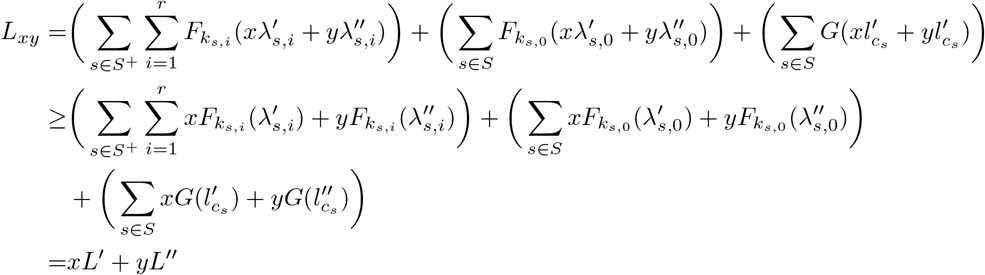

Where equality holds if and only if 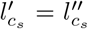 for all *s* and 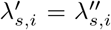 for all *s* ∈ *S*^+^ and *i* due to the strict concavity of *F* and *G* given by lemma 1. Then it must be the case that 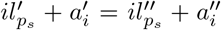 for all *s* and We defined 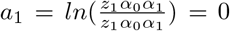, thus 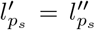 for all *s* ∈ *S*^+^. Since there must exist an *s* such that its read vector has no zero entries by assumption, that also means that 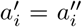 for all *i* as well. Thus, the two parameter sets are the same. This is a contradiction, since we supposed that the two parameter sets are different, so it must be the case that 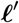 is strictly concave.

Supposing that the maximum likelihood estimate is not unbounded, it must be that case that it is unique since 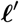 is strictly concave. It remains to show that the estimate is not unbounded.

### Lemma 3.

*For all b, the set of parameters that achieves* 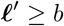 *is bounded*.

*Proof.* Each of the terms in equation 24 is bounded from above by lemma 1, and drops without bound as the magnitude of their arguments tend to infinity. Therefore, the last set of terms given by 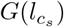 imply that to attain some *b*, the magnitude of 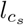 cannot go beyond some value, since 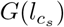 approaches in either direction, and the negativity cannot be compensated for by other terms.

By the same argument, 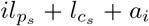 must be similarly bounded for all *s* and 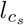 where *k*_*s,i*_ > 0. Since *a*_1_ = 0 and *l_c_s* are bounded for all *s*, that means 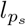 are similarly bounded for all *s* where *k*_*s,*1_ > 0. Since there exists a read vector with no zero entries by assumption, that then implies that *a*_*i*_ are similarly bounded for all *i*, which in turn implies that 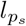 are bounded for all *s*. Thus, the region of parameters for which 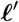 attains some give *b* is bounded.

If we let *b* be the maximum likelihood, then the above lemma shows that the parameters that attain it cannot be unbounded. Thus, RSR values exist and are uniquely determined given that there exist at least one read vector with no zero entries. Since the objective is strictly concave and we can bound the region to consider after given an initial guess of the maximum likelihood, we can apply convex optimization techniques to obtain RSR values.

## B Proof of Theorem 2 (Basis features for unique decompositions)

The set of all functions on sequences *Σ^n^* → ℝ form a vector space, with vector addition defined as *f* + *g* = *h* where *h*(*s*) = *f* (*s*) + *g*(*s*) for all *s* ∈ *Σ*^*n*^, and scalar multiplication defined as *cf* = *h* where *h*(*s*) = *cf* (*s*) for scalar *c* and all *s* ∈ *Σ*^*n*^. This space is spanned by the set of functions that evaluate to 1 on some given sequence and 0 to all other sequences. These functions are linearly independent, so this vector space has a finite dimension of |*Σ^n^*|.

Sequence features are functions on sequences, so they are elements of this space. It follows from definition that order *k* functions form a subspace which is spanned by the sequence features of order *k* or less.

Let INV be the fixed invariant symbol. We say that a first order sequence feature contains INV if it is of the form 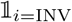, and more generally that a sequence feature contains INV if any of its first order factors contain INV.

### Lemma 4.

*Let B_k_ be a set of vectors spanning for the space of order k functions. Then B_k_* ⋃ *U is a set spanning the space of order k* + 1 *functions, where U is the set of order k sequence features that do not contain* INV.

*Proof.* It is enough to show that we can construct all sequence features of order *k* + 1 using a linear combination of vectors in *B_k_ ⋃ U*. We will construct them inductively. In the base case, we have all the sequence features of order *k* + 1 with no factor containing INV.

As an induction hypothesis, suppose that we can construct all sequence features of order *k* + 1 that contains at most *a* first order factors that contain INV. We want to be able to construct any order *k* + 1 sequence feature that contains *a* + 1 factors that contain INV. Let *b* be such a sequence feature.

Let *b*’ be an order *k* sequence feature such that *b* ⊆ *b*’, and that *b* contains exactly *a* factors containing INV. We can then express *b* as:

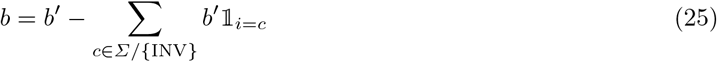

Where *i* is the position occupied by *b* but not by *b*’. *b*’ is an order *k* sequence feature, so it is contained in *B*_*k*_ and we can construct it. All 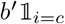 has at most *a* factors that contain INV, so we can construct them by the induction hypothesis. Therefore, an arbitrary order *k* + 1 sequence feature with *a* + 1 first order factors that contain INV can be expressed as a linear combination of sequences features of order *k* + 1 or less that contain at most *a* factors that contain INV.

Carrying on the induction until *k* + 1 shows that any sequence feature of order *k* + 1 can be constructed, which completes the proof.

Therefore, we can express any function by only using sequence features that do not contain INV. For uniqueness, we need to show that these vectors are linearly independent.

Suppose for contradiction that they are not. That means the number of sequence features that don’t contain INV must be strictly larger than the dimension of the space, since some of the sequence features are linearly dependent, and the full set must span the entire space. The number of sequence features that don’t contain INV is:

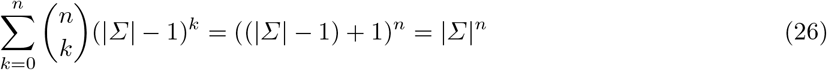

This should be strictly larger than the dimension of the vector space, which we showed to be |*Σ|^n^*. It isn’t, which leads to a contradiction. Therefore the vectors in our construction must all be linearly independent.

Therefore, for each function there is exactly one linear combination of sequence features without INV that equals it.

Finally, to see that if we have an order *k* function we won’t have any coefficients greater for sequence features greater than order *k*, note that lemma 4 shows that the set of order *k* functions is already spanned by the sequence features of order *k* that don’t contain INV, so they are sufficient to express any order *k* function.

## C Proof of Theorem 3 (Soundness and parsimony of the canonical decomposition)

The uniqueness of the canonical decomposition follows from its definition. To see that the canonical decomposition is a decomposition, note that by rearranging equation 14, the canonical decomposition *C* of a function *f* has the following property for any for any order *k* sequence feature *q*:

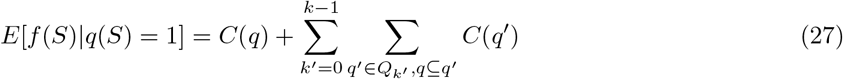

Where again *S* denotes a random sequence distributed uniformly over *Σ^n^*. If *q* is an order *n* sequence feature corresponding to sequence *s*, then the left hand side becomes *E*[*f* (*S*)|*S* = *s*] = *f* (*s*) and the right hand side becomes precisely the function of *C* evaluated on *s*. Therefore, the function of *C* is *f*, so *C* is a decomposition of *f*.

We now show that it is also maximally parsimonious. We showed in appendix B that *Σ^n^* → ℝ has a vector space structure. We now define an inner product on that space, where 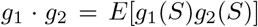. It follows that we can define the squared norm as 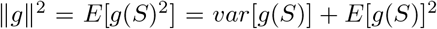. We note that for any sequence feature *q*, we have *g* · *q* ∝ *E*[*g*(*S*)|*q*(*S*) = 1].

Consider a decomposition *P* of *f*. We denote the function of *P* [*k*] as 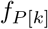. Suppose *P* has the following orthogonality property: for each *k* ∈ [*n*]^+^, 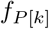 is orthogonal to the space spanned by sequence features of order *k* − 1.

### Lemma 5.

*P is at least as parsimonious as any other decomposition of f*.

*Proof.* Let *P*’ be any other decomposition of *f*. Denote the function of the *k*th component *P*’[*k*] as 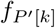. Suppose that 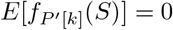 for all *k* ∈ [*n*]+. For any decomposition, there is an equivalent decomposition with this property that expresses the same function, and where the set of decompositions that are at least as parsimonious as it and the set of decompositions that it is at least as parsimonious as remain unchanged. This is achieved by shifting coefficients into the order 0 component throughout every other component of the decomposition, which does not affect the variance of any component and therefore does not affect parsimony. Therefore, the supposition that 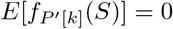 for all *k* ∈ [*n*]+ can be made without loss of generality.

Let *k* be the largest value for which 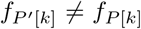. If no such *k* exists, then we are done, since we must have *P* = *P*’, and so *P* is at least as parsimonious as *P*’. Therefore, assume such a *k* exists. Note that this *k* cannot be zero if it does, since that would mean the functions of the two decompositions differ by a constant, which implies that they express different functions.

We claim that 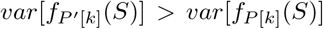, which would be sufficient to prove the lemma. We have 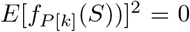, since *k* = 0 and 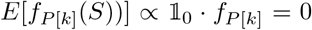 from the orthogonality property of *P*. We have 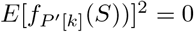 by assumption. Therefore, it is sufficient to show that 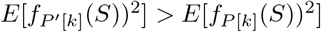, or alternatively 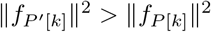.

We claim that 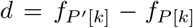 is an order *k* − 1 function. This is because any component of 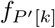 orthogonal to the space of order *k* − 1 functions must equal the component of 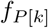 orthogonal to that space. That in turn is because the difference between the orthogonal components of the functions cannot be compensated for by higher order components since they are all equal, and cannot be compensated for by lower order components since they cannot express functions orthogonal to them. Therefore, in order for the decompositions to express the same function it is necessary for the components of 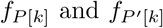 that are orthogonal to lower order functions to be equal.

Then 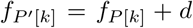, where 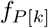 and *d* are orthogonal since 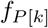 is orthogonal to any order *k* − 1 function. By the Pythagorean theorem on arbitrary inner product spaces, we have 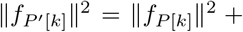 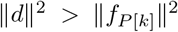. This proves our claim that 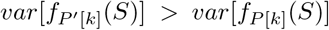. Therefore, *P* is at least as parsimonious as *P*’, which completes the proof.

We now show that the canonical decomposition has this orthogonality property held by *P*, which will then suffice to prove that it is at least as parsimonious as any other decomposition of *f*.

First, given a set of positions we define a positional variable of a decomposition as the sum of all the terms in a decomposition that occupy exactly those positions. The order of a positional variable is the number of positions occupied. The *k*th order component of a decomposition is the sum of all its *k*th order positional variables.

### Lemma 6.

*Given a canonical decomposition C, if V is a kth order positional variable of C then V is orthogonal to all functions of order k* − 1.

*Proof.* We will prove this by induction. In the base case, we let *k* = 0 where there exists no function of order −1, so the lemma is trivially true.

Suppose that the lemma holds for order up to *k* − 1. We will show that it holds for order *k* as well.

Consider a positional variable *V* of order *k* that occupies a set of positions *X*. Let *V*_*s*_ be the set of sequence features that occupy the position *V* does. Then 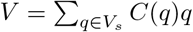. Let *q*’ be any sequence feature of order less than *k*. Let *q*’’ be the same sequence feature, but with only the first order factors that occupy locations in *X*. Define the set of sequence features 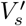 as

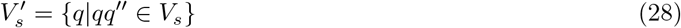

By total expectation, we have:

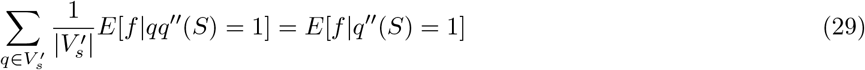

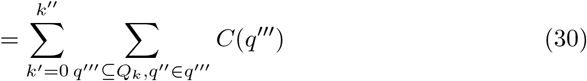

Where the last equality follows from equation 27, and *k*’’ is the order of *q*’’. Note that since *q*’ has order less than *k*, so must *q*’’, therefore *k*’’ < *k*. By rearranging and applying equation 27 once again we obtain:

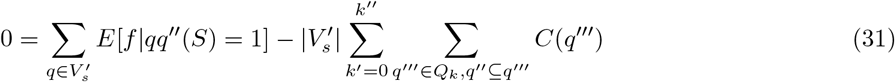

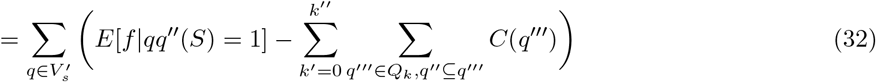

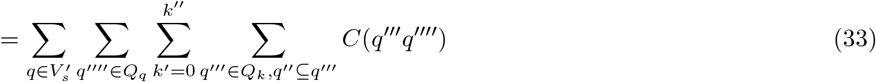

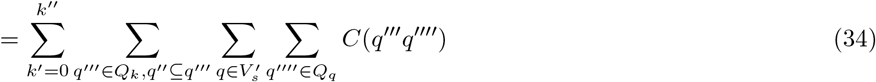

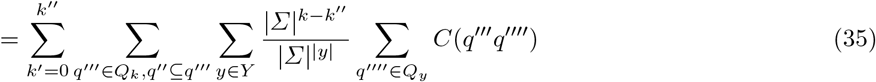

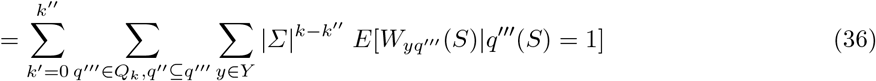

Where *Q*_*q*_ denotes the set of sequence features *x* that satisfy *q* ⊆ *x* and 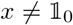. *Y* denotes the power set of the set of positions that are occupied by variables in 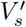 minus the empty set. *Q*_*y*_ denotes the set of sequence features that occupy exactly the positions *y*. 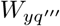 denotes a positional variable that occupies the positions y and the positions occupied by *q*’’.

By our induction hypothesis, for all 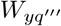 that are of order less than *k* we have 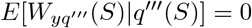. Therefore, the remaining term is:

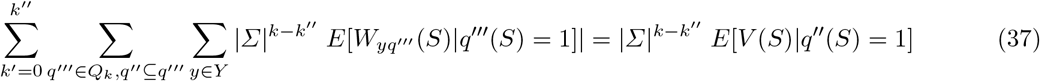

Which gives us:

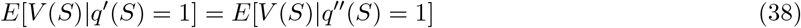

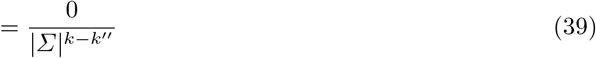

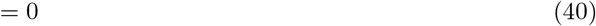

Since *E*[*V* (*S*)| *q*’(*S*) = 1] ∝ *V* · *q*’, the positional variable *V* is orthogonal to any arbitrary lower order sequence feature *q*’. Since all lower order functions can be expressed as a linear combinations of lower order sequence features, *V* is orthogonal to all lower order functions. Therefore all positional variables of order *k* are orthogonal to all functions of order *k* − 1.

Carrying on the induction until *n* proves the lemma for all positional variables.

Since the function of a component of order *k* can be expressed as a sum of positional variables of order *k*, the canonical decomposition has the orthogonality property that we assumed *P* does. Therefore, it is at least as parsimonious as any other decomposition by lemma 5.

## D Running Peptide Display in Silico

To sample from the theoretical distribution of peptides after *r* rounds of peptide display, we note that the ratio of concentrations between 9-mer *s* and 9-mer *s*’ can be obtained by rearranging equations 2 and 6 to obtain the following:

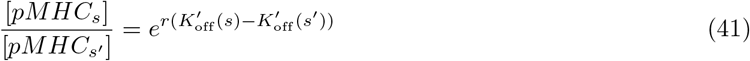

Where *r* is the number of rounds. We can then substitute 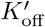 with 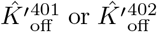 to obtain the values given by our model of reaction kinetics. This allows us to obtain an unnormalized marginal distribution of residues at a given position after a given number of rounds, which we can then normalize to obtain the marginal distribution of residues at that position, conditioned on the identities of the other residues.

We perform Gibbs sampling over this distribution, where we consider each sample to be a vector of 9 residues. We initialize with a random sequence selected from the uniform distribution, run for a burn-in period of 10^5^ iterations, and then collect samples over the following 10^6^ iterations. We then report the expected value of first order sequence features approximated from the samples, which we then use to construct sequence logos for comparison against the experimentally derived distributions. We compare the distributions after each round in figures 4 and 5. We note that the divergence in distributions in early rounds may be partially attributed to the fact that the randomized library for yeast display does not uniformly contain all 20^9^ sequences.

**Fig. 4.**
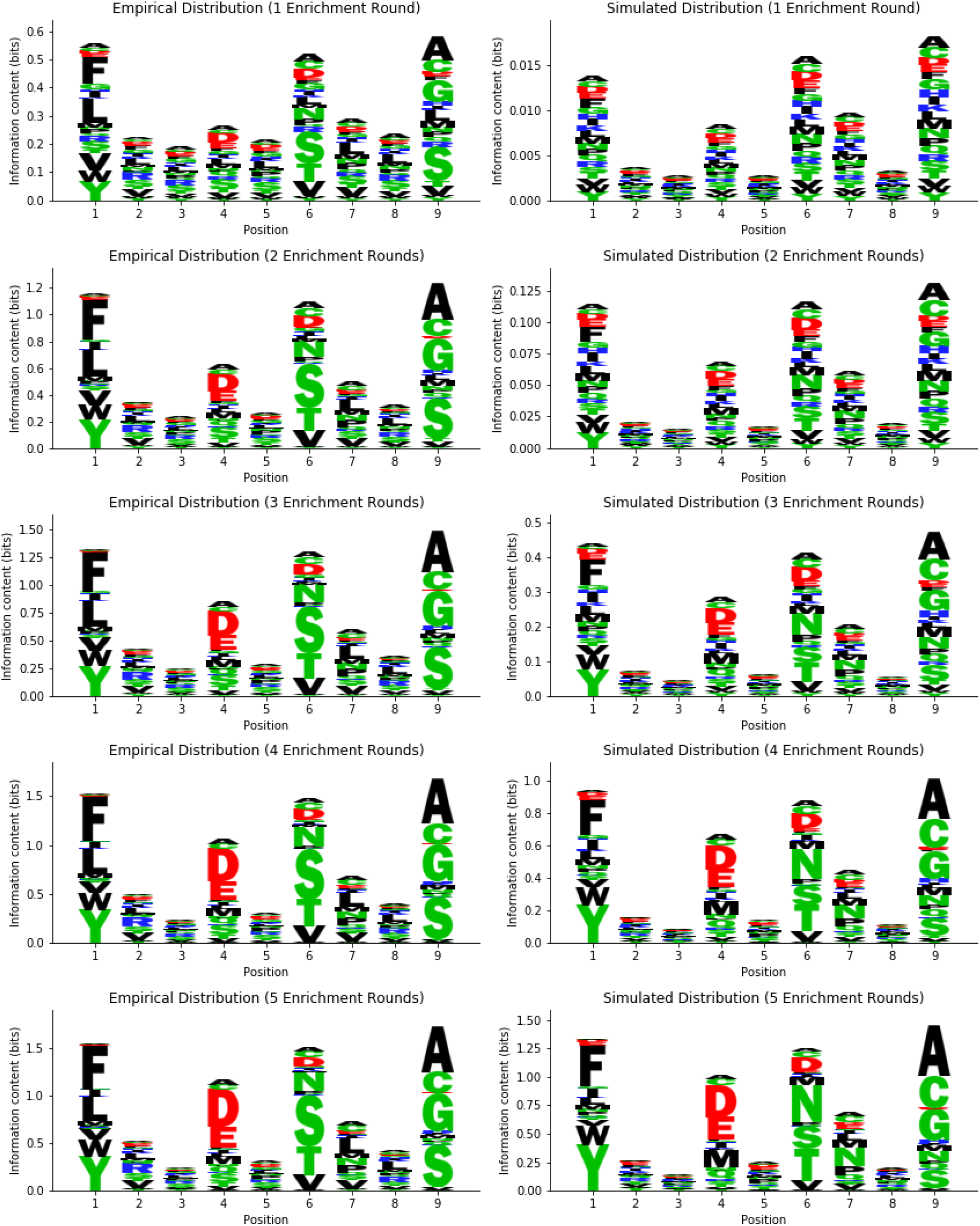
Empirical progression of the distribution against the predicted progression for DR401.

**Fig. 5.**
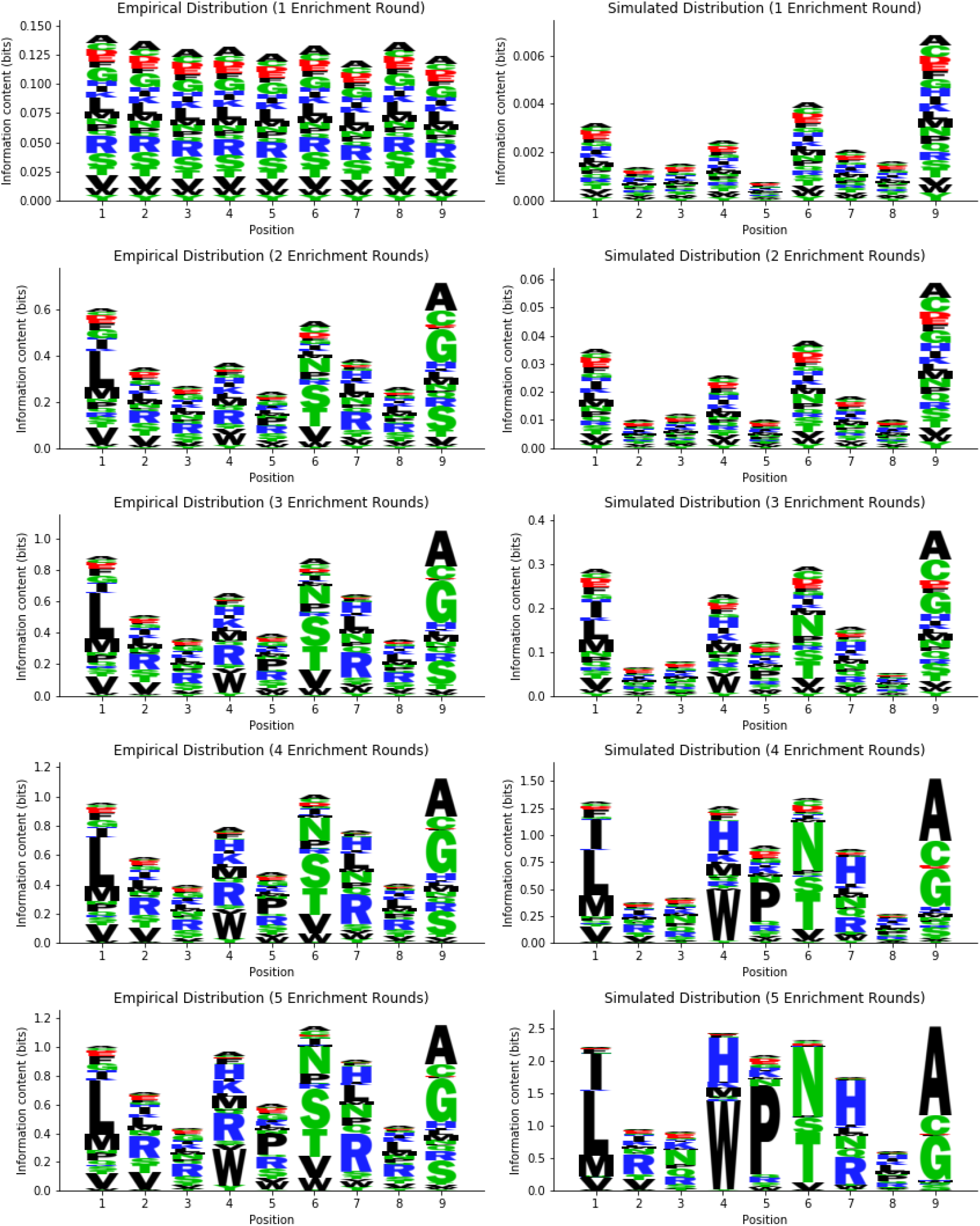
Empirical progression of the distribution against the predicted progression for DR402.

